# Reduction of Neuroinflammation and Seizures in a Mouse Model of CLN1 Batten Disease using the Small Molecule Enzyme Mimetic, N-Tert-Butyl Hydroxylamine

**DOI:** 10.1101/2024.05.01.591955

**Authors:** Zach Fyke, Rachel Johansson, Anna I. Scott, Devin Wiley, Daniel Chelsky, Joseph D. Zak, Nader Al Nakouzi, Kevin P. Koster, Akira Yoshii

## Abstract

Infantile neuronal ceroid lipofuscinosis (CLN1 Batten Disease) is a devastating pediatric lysosomal storage disease caused by pathogenic variants in the *CLN1* gene, which encodes the depalmitoylation enzyme, palmitoyl-protein thioesterase 1 (PPT1). CLN1 patients present with visual deterioration, psychomotor dysfunction, and recurrent seizures until neurodegeneration results in death, typically before fifteen years of age. Histopathological features of CLN1 include aggregation of lysosomal autofluorescent storage material (AFSM), as well as profound gliosis. The current management of CLN1 is relegated to palliative care. Here, we examine the therapeutic potential of a small molecule PPT1 mimetic, N-tert-butyl hydroxylamine (NtBuHA), in a *Cln1^−/−^* mouse model. Treatment with NtBuHA reduced AFSM accumulation both in vitro and in vivo. Importantly, NtBuHA treatment in *Cln1*^−/−^ mice reduced neuroinflammation, mitigated epileptic episodes, and normalized motor function. Live cell imaging of *Cln1*^−/−^ primary cortical neurons treated with NtBuHA partially rescued aberrant synaptic calcium dynamics, suggesting a potential mechanism contributing to the therapeutic effects of NtBuHA in vivo. Taken together, our findings provide supporting evidence for NtBuHA as a potential treatment for CLN1 Batten Disease.

## 1. Introduction

The neuronal ceroid lipofuscinoses (NCLs), historically identified as Batten Disease, represent a spectrum of autosomal recessively inherited neurodegenerative diseases caused by pathogenic changes in one of 14 genes identified to date [1–3]. Infantile NCL (aka CLN1, Batten Disease type 1) is a particularly devastating subtype [3]. Symptom onset in CLN1 typically presents around six months of age with progressive blindness, psychomotor dysfunction, and recurrent seizures, ultimately culminating in fatality by 15 years of age [4–10].

Seizures in patients with CLN1 may present before a diagnosis is made and are often present very early in the disease course [8]. Seizures can be as frequent as multiple times a day with types varying from focal to primary generalized seizures [8]. Seizures in CLN1 Batten Disease are often debilitating for patients and challenging for caregivers. Current management of seizures include existing anti-seizure medications with varying efficacy. Depending on seizure type, duration, and age of onset, the neurocognitive effects can range from synaptic-circuit reorganization to neuronal cell death [11]. More effective seizure control earlier in the disease course might improve quality of life for patients and caregivers, as well as reduce the long-term harmful effects of repeat seizures on neuronal function.

CLN1 is caused by a pathogenic mutation of the depalmitoylation enzyme, palmitoyl-protein thioesterase 1 (PPT1). Protein palmitoylation (S-acylation) is a reversible post-translational modification linking palmitate, a 16-carbon fatty acid, to proteins [12]. Palmitoylation dynamics influence multiple cellular functions and are critical for regulating protein trafficking and degradation [12–14]. Palmitoylation is orchestrated by more than twenty palmitoyl acyltransferases (PATs), frequently localized to the Golgi apparatus, where they catalyze covalent bonding of palmitate to target proteins [15,16]. Depalmitoylation is carried out by only a few enzymes, including PPT1, which localizes to several cellular compartments but chiefly the lysosome [14,17,18]. Accordingly, autofluorescent storage material (AFSM) consists of accumulated palmitoylated proteins in lysosomes and is a histopathological hallmark in CLN1. AFSM is a shared microscopic finding of NCLs and was described before the identification of disease-causing genes [3,6,7].

Among all the NCLs, there is currently only one clinically approved, disease-modifying treatment that is exclusive to CLN2 [19]. For CLN1, disease-modifying therapeutics are not commercially available. Current treatments focus on symptom management and palliative care measures [8]. While gene and enzyme replacement therapies successfully mitigate disease symptomology in CLN1 mouse models [20–24], enthusiasm for elevating these therapies to human clinical trials hinges on optimizing their delivery and expression, as well as their safety profile [25]. An adjunctive strategy to treat CLN1 is small molecule therapeutics, particularly those that mimic the endogenous function of PPT1. In previous work, Sarkar and colleagues demonstrated that the hydroxylamine derivative N-tert-butyl hydroxylamine (NtBuHA) cleaved the thioester linkage of post-translational palmitoyl modifications, reduced histopathological markers of CLN1, improved motor function, and increased lifespan in *Cln1^−/−^*mice [26].

Here, we tested the therapeutic efficacy of PPT1-mimetic supplementation in *Cln1^−/−^* mice [27] using a novel formulation of NtBuHA that improves its pharmacokinetic profile, called CIRC827. We discovered that this novel formulation of NtBuHA reduced AFSM accumulation both in vitro and in vivo, improved primary neuronal calcium dynamics, attenuated neuroinflammation, mitigated seizures, and rescued motor function in *Cln1^−/−^* mice. These results confirm that small molecule PPT1-mimetic treatment is a potential therapeutic approach for CLN1 and provides novel mechanistic insight into its beneficial effects.

## 2. Materials and methods

### 2.1 Animals, genotyping, group allocation, and data handling

Animal procedures, genotyping, and data handling were performed as in Koster et al [28,29]. Ppt1+/-(heterozygous) mice were obtained from Jackson Laboratory and maintained on 12 hours light/dark cycle with food and water ad libitum. Animals were genotyped in-house as first demonstrated [27] and used for experiments at specified multiple disease-relevant time points: postnatal day 60 (P60) and P180. After verifying the genotype of animals via toe-clip before P10, WT and *Cln1^−/−^* mice were enrolled to treatment conditions at P30 including vehicle (H2O), 250μM, 1mM, and 4mM NtBuHA for either 30 days (P60) or 150 days (P180) respectively. Data was either analyzed by lab members blinded to condition or randomized prior to analysis by ZAF and JDZ. All animal procedures were performed in accordance with the guidelines of the University of Illinois of Chicago Institutional Animal Care and Use Committee (Protocol# 21-075).

### 2.2 NtBuHA Drug solubilization and administration

NtBuHA (Circumvent Pharmaceuticals, see *Key Resources Table*) was received as 40-gram aliquots in amber glass jars wrapped in parafilm and kept at room temperature out of direct sunlight. Since NtBuHA is hydrophilic, the compound was solubilized via serial dilutions ranging from 0.1μM to 1mM NtBuHA in PBS pH 7.4 for the treatment of primary cortical neuron culture. For *ad libitum* dosing to wild-type (WT) and *Cln1^−/−^* animals, NtBuHA was prepared in standard animal care facility water bottles at a concentration of 250μM, 1mM, and 4mM. Fresh drug aliquots were administered by lab members weekly, with animals’ weights monitored from P30-60 for potential side effects of drug treatment.

### 2.3 Primary cortical neuron culture and transfection

Primary cortical neuron culture was conducted as described [28,29]. Briefly, timed pregnancies from *Cln1^-/+^*dams were grown until embryonic day 15.5 (E15.5). For live-cell imaging experiments, cells were counted and then plated at 350,000-400,000 cells/dish on 35mm cell culture dishes containing poly-D-lysine/laminin-coated coverslips (Cellvis). Cells were grown until DIV7 (day in vitro), and then were transfected as described in Koster et al [28,29]. In summary, GCaMP_3_ (4μg DNA/dish) was combined in a transfection construct containing both Lipofectamine 2000 (2μg/dish) and Neurobasal plus B27 neuronal culture media. Transfection solution was applied to the primary neurons which were incubated at 37°C for 45 minutes. Neurons recovered until DIV14, at which point *Cln1^−/−^* samples were treated with 100μM NtBuHA every 48 hours from DIV12-18. (See *Key Resources Table*)

### 2.4 Immunocytochemistry and microscopy imaging and analysis

For immunocytochemistry (ICC) experiments, primary neurons were dissociated, cells were counted then plated at 150,000-180,000 cells/well in 24-well plates containing poly-D-lysine/laminin-coated coverslips. ICC neuronal culture was grown until DIV7, then treatment was administered via a 1:1,000 dilution of NtBuHA at indicated concentrations every 48 hours until DIV 18. Finally, cells were fixed in 4% paraformaldehyde (PFA) solution. Immunostaining was performed as described previously [29]. In short, neurons were stained for microtubule-associated protein 2 (Anti-MAP2, Rabbit; Millipore-Sigma; RRID: AB_5622). Following primary antibody incubation, coverslips were then incubated in the fluorophore-linked secondary antibody (Alexa Fluor 633 Goat anti-Rabbit; Invitrogen Antibodies; RRID: AB_2535731). Coverslips were mounted on SuperFrost Plus slides in DAPI Vectamount medium. To investigate AFSM accumulation after NtBuHA treatment, primary neurons were grown until DIV7, then treated with 1:1,000 dilutions of either vehicle (PBS) or 0.1-1mM NtBuHA every 48 hours until DIV19. After the final treatment, cells were washed in PBS before fixation in 4% PFA in PBS.

For in vitro AFSM particle detection, confocal imaging was performed using the LSM710 confocal microscope with Zen Black imaging software. Z-stack max projection images of monolayer primary neurons (for a total of 10 microns of z-plane depth to properly encompass the entirety of the cell’s soma) were collected. Next, the open-source image analysis software FIJI was used to create max-projection compressions of all imaging channels including DAPI (405nm), AFSM (555nm), and MAP2 (633nm). Next, image channels were separated, a single neuron was selected, and the cell soma was encircled as a region of interest (ROI). Then, the corresponding nucleus was cleared across channels to isolate somatic fractions containing AFSM. The default threshold algorithm was used to generate a binary mask of AFSM puncta. The number of puncta, total area, and percent area were derived using the ‘analyze particles’ tool (>2^2^ pixels). These data were then input to Prism Graph Pad 9 for graphs and statistics. (See *Key Resources Table*).

### 2.5 Transcardial perfusion and immunohistochemistry

Transcardial perfusion, immunohistochemistry, and image analysis were performed as described previously [28,29]. Briefly, untreated and NtBuHA treated *Cln1^−/−^* and WT mice were anesthetized with isoflurane and transcardially perfused with PBS followed by 4% PFA in PBS. Brains were removed and fixed overnight at 4°C in PFA, then transferred to 30% sucrose in PBS before sectioning. 100μm coronal sections were made in cold PBS using a Vibratome 1000 (Technical Products International, St. Louis, MO). Four serial sections per well were stored in a cryoprotectant solution (30% glycerol, 30% ethylene glycol in PBS) at −20°C until immunohistochemistry was performed. 3-4 brain slices per animal were isolated from primary somatosensory cortex (S1) and primary visual cortex (V1) and were first incubated in PBS before undergoing permeabilization. Next, samples underwent antigen retrieval by heating in Tris-EDTA. Brain sections were blocked (TBS + 0.1% Triton X-100, 4% BSA, and 5% normal goat serum) before being incubated in rabbit anti-Iba1 (Anti Iba1, Rabbit; Fujifilm; Code No. 019-19741) for 48 hours. After 4 PBS washes of 10 minutes each, tissue was incubated in Alexa Fluor 633 (Goat anti-Rabbit; Invitrogen Antibodies; RRID: AB_2535731). After washing, the tissue was incubated in GFAP antibody (Anti-GFAP, Goat; Abcam; RRID: AB_53554) for 48 hours. Then, the slices were incubated in Alexa Fluor 488 (Donkey Anti-Goat IgG; Jackson ImmunoResearch Laboratories; RRID: AB_2336933). Finally, two slices from S1 and two slices from V1 were mounted on SuperFrost Plus slides in DAPI Vectamount medium. (See *Key Resources Table*)

### 2.6 In Vivo Histology Analysis

Two z-stack images from L2/3 of S1 and V1 from each animal were acquired using a Zeiss LSM710 confocal laser scanning microscope at 63x magnification. All sections were imaged using identical parameters. Quantification of AFSM was performed by creating a max projection z-stack compression (20 micron/ 1 image per micron) and subsequent background subtraction and automated images thresholding in FIJI (See *Key Resources Table*), generating an 8-bit binary mask of AFSM-positive pixels (>5^2^ pixels). The identical threshold was applied to each image (Default threshold algorithm). The number of AFSM particles above threshold, as well as percent area occupied by AFSM puncta, the average puncta size, and total AFSM area was then calculated using the “analyze particles” tool. GFAP histology was performed as described for AFSM particles, with the threshold for astrocyte arbors set at >15^2^ pixels. Individual GFAP astrocyte counts were made by an analyst, blinded to treatment condition, and were described as a fully stained cell body along with 4 directly connected processes. IBA1 microglial Scholl analysis was completed as previously described [29]. Briefly, 20 μm z-stack compressions were made in the 633nm confocal imaging channel. Three microglia per image, 5 images from S1 and 5 images from V1 were isolated and thresholded using the ‘default’ algorithm. An ROI was set in the center of the cell soma, to which we applied the neuroanatomy plug-in ‘skeletonize’ to trace microglial ramifications. Next, we used the Scholl Analysis plug-in, setting the concentric ring distance at 2 μm step size, with the first concentric ring starting at 5μm from the cell soma and expanding to 70 μm. The number of microglial ramifications that intersect with each ring was graphed, with the corresponding distance from the cell soma indicated. For all IHC experiments, three animals per condition were used for analysis.

### 2.7 Live Cell Imaging

Live cell imaging was performed as previously described [29]. At DIV8, primary cortical neuron cultures from WT and *Cln1^−/−^* cultures were transfected to express GCaMP_3_ as described above. Then, cultures were maintained until DIV12 where each dish received either vehicle (PBS) or 100μm NtBuHA via 1:1,000 dilution into cell culture media. Treatment was administered once on DIV12 and again on DIV14, two hours prior to live cell imaging. Next, 4-minute videos of a single transfected neuron were taken at 10x magnification using the LSM 710 confocal microscope at a rate of 4 frames/ second.

### 2.8 Calcium Imaging Analysis

Images were exported as raw TIFF files for processing using custom and available MATLAB scripts. Prior to processing, images were binned at 2 × 2 by averaging pixel values in each bin. Non-rigid motion artifact compensation and denoising were then performed using NoRMcorre [30]. The CaImAn CNMF pipeline [31] was used for ROI selection (See *Key Resources Table*). ROIs were further filtered by size and shape to remove merged cells. The analysis pipeline was optimized for in vitro imaging [32] and for the selection of subcellular structures [33]. Calcium transients were identified by local maxima and included if the peak amplitude exceeded 3.5 standard deviations of a noise distribution obtained from baseline samples. Instantaneous event frequencies were calculated by considering the rate of spontaneous calcium events across all synaptic sites associated with a single neuron and interleaving event timestamps.

### 2.9 EEG transmitter implantation & Data Acquisition

EEG transmitter implantation was performed as recently described [34]. Briefly, 6.5-month-old WT and *Cln1^−/−^* animals were anesthetized using 1.5-3% isoflurane and then head-fixed in the stereotaxic apparatus. Hair was removed from the scalp of the animal, and the skin was sterilized with ethanol wipes and iodine swabs. A 1.5 to 2 cm incision was made along the midline of the skull just anterior to bregma to the rostral border of the trapezius muscles. Then, an ETA-F10 wireless radio-telemetric transmitter weighing 1.6 grams (ETA-F10, Data Sciences International, St. Paul, MN) was inserted into a subcutaneous pocket, made using blunt scissors above the left hindlimb. Two biopotential leads were fixed unilaterally and subdurally above the motor cortex (M1) and visual cortex (V1) by screws. Each lead was wrapped securely around a single screw. The leads and screws were secured by covering with dental cement (See *Key Resources Table*). Once the cement was dry, the incision was closed using 4-0 monofilament suture with a simple interrupted pattern. After the suture was completed, antibiotic ointment was liberally applied to the incision prior to the animal’s recovery on a heating pad.

### 2.10 Telemetric recording of Unilateral EEG Activity

After recovery from surgery, mice were moved to recording cages to acclimatize. At the time of recording, telemetric transmitters were magnetically activated. EEG data was collected as described previously [35]. The cages were mounted on a DSI telemetry receiver (RPC-1), connected to a DSI Data Exchange Matrix (MX1) and subsequently to an acquisition computer. EEG recordings were conducted 24-36 hours post-implantation using Ponema (Ver 5.20). Once 24-hour EEG recordings were complete, telemetry units were inactivated, and animals were euthanized at 72 hours.

### 2.11 EEG Data processing and analysis

EEG traces were uploaded as .RAW files and imported into DSI’s analysis software (Neuroscore Ver 3.1). For data analysis, the dynamic spike detection threshold was manually configured to identify epileptiform spikes >5x above baseline EEG signal. The spike detection algorithm was used to generate seizure reports for each animal by an experimenter, blinded to the treatment condition. Data were generated and analyzed as described [34], using a manual analysis paradigm. Additionally, a subset of EEG recordings included video corresponding to time-stamped EEG traces for a quantitative confirmation of severe convulsive epileptic episodes from both untreated and NtBuHA-treated *Cln1^−/−^* mice (**Supplementary Video 1-2**). Prism 9 and Biorender were used to generate graphs and statistics and illustrations.

### 2.12 Rotarod Testing

Single-speed rotarod testing was completed in Wild Type and *Cln1^−/−^*mice at 2 months, 4 months, and 6 months of age. For each test, animals were acclimatized to the testing room for one hour prior to training. Next, animals were placed on an ENV-577 with the speed set to 16 RPM for 1 minute. Two training trials of one minute were completed, with a minimum of 10 minutes break between training rounds. After training was complete, the animals were placed on the rotarod at 16 RPM for 3 minutes, with the latency to fall recorded by an experimenter, blinded to treatment condition. At the cessation of the trial, three testing round scores were averaged to generate the average latency to fall. Prism 9 was used to generate graphs and statistics.

## 3. Results

### 3.1 NtBuHA administration reduces AFSM in Cln1^−/−^ primary cortical mouse neurons

First, we tested whether NtBuHA could reduce the aggregation of AFSM in primary cortical neurons derived from embryonic *Cln1^−/−^* mice. Wild-type (WT) and *Cln1^−/−^* mouse primary neurons were treated with either vehicle (PBS) or NtBuHA every 48h for 12 day (**Supplemental Fig. 1A, 1B**). Following the final treatment, neurons were fixed, immunostained for MAP2 and analyzed for AFSM accumulation (**Fig. 1A**). Automated particle analysis of AFSM in neuronal soma showed that, as expected, WT neurons did not accumulate AFSM, while untreated *Cln1^−/−^* primary neurons typically contained multiple AFSM puncta (**Fig. 1A**). Importantly, after treatment with NtBuHA, we observed a dose-dependent reduction in the number of AFSM puncta compared to untreated *Cln1^−/−^* neurons (**Fig. 1B**). Similarly, the percent area of AFSM deposits within the soma of NtBuHA-treated *Cln1*^-/*-*^ primary neurons (**Fig. 1C**) and size of puncta (**Supplemental Fig. 1C**) were also reduced.

**Figure 1:**
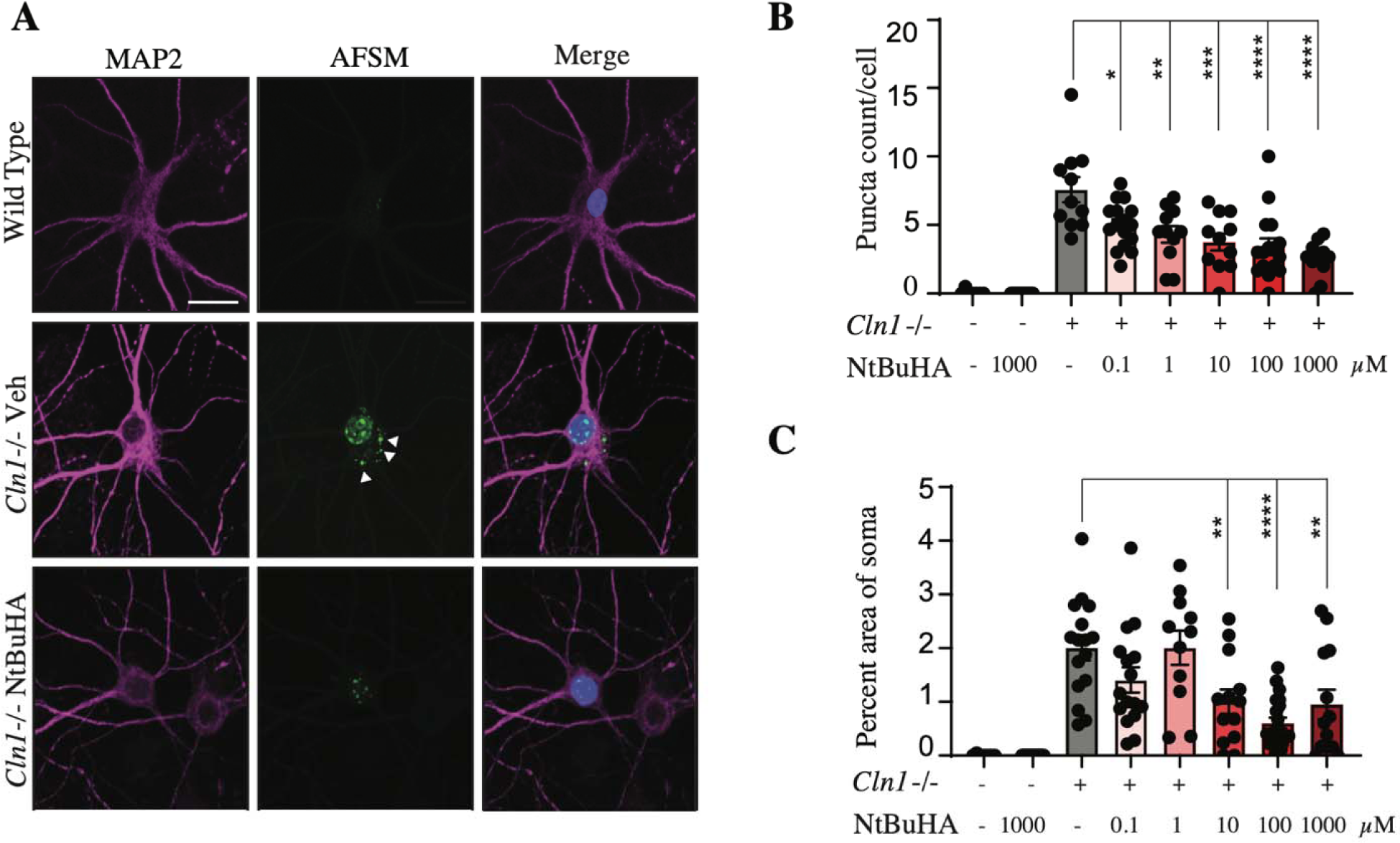
NtBuHA Reduces AFSM Accumulation in *Cln1^−/−^* Neuron Culture. **(A)** Representative images of WT, *Cln1^−/−^*, and NtBuHA-treated *Cln1^−/−^* neurons immunostained for MAP2 and DAPI and demonstrating the presence of AFSM deposits in neuronal somata (arrowheads). Scale=25_μ_m. (**B)** Quantification of AFSM puncta count per cell, *Cln1^−/−^* vehicle-treated=7.488 particles, n=29; comparison to *Cln1^−/−^* 0.1_μ_M NtBuHA=5.157 particles, n=28, P=0.0358; comparison to *Cln1*^−/−^ 1_μ_M NtBuHA=4.511 particles, n=25, P=0.0042; comparison to *Cln1*^−/−^ 10_μ_M NtBuHA=4.071 particles, n=25, P=0.0002; comparison to *Cln1*^−/−^ 100_μ_ NtBuHA=3.071 particles, n=35, P<0.0001; comparison to *Cln1*^−/−^ 1_μ_M NtBuHA=2.711 particles, n=28, P<0.0001, Šídák’s multiple comparisons test **(C)** percent area of AFSM puncta per cell soma, *Cln1*^−/−^ vehicle-treated=2.062 percent of soma, n=29; comparison to *Cln1*^−/−^ 10_μ_M NtBuHA=0.8055 percent of soma, n=25, P=0.0012; comparison to *Cln1*^−/−^ 100_μ_M NtBuHA=0.5546, n=35, P=0.0089; comparison to *Cln1*^−/−^ 1m NtBuHA=0.8754, percent of soma, n=28, P=0.0095, Šídák’s multiple comparisons test.

### 3.2 Treatment with NtBuHA reduces AFSM accumulation and attenuates neuroinflammation in Cln1^−/−^ mice

To validate and extend our in vitro findings, we tested the effect of early therapeutic supplementation of NtBuHA in *Cln1^−/−^* mice. Beginning at postnatal day 30 (P30), *Cln1^−/−^* mice were given free access to either drinking water or water supplemented with NtBuHA. We then analyzed AFSM accumulation in the primary somatosensory (S1) and visual cortices (V1) of treated and untreated *Cln1^−/−^* animals at 2 (P60) and 6 months (P180) [36]. As demonstrated previously [28], AFSM accumulation was modest by P60 and robust by P180 in untreated *Cln1^−/−^*mice. Both the AFSM puncta count (**Fig. 2A, 2B**) and percent of image area occupied by AFSM (**Fig. 2C**) were significantly diminished in NtBuHA-treated *Cln1^−/−^*mice at 2 months. Further, while NtBuHA supplementation did not reduce AFSM puncta count at 6 months, it did diminish the percent area of AFSM (**Fig. 2B, 2C**).

**Figure 2.**
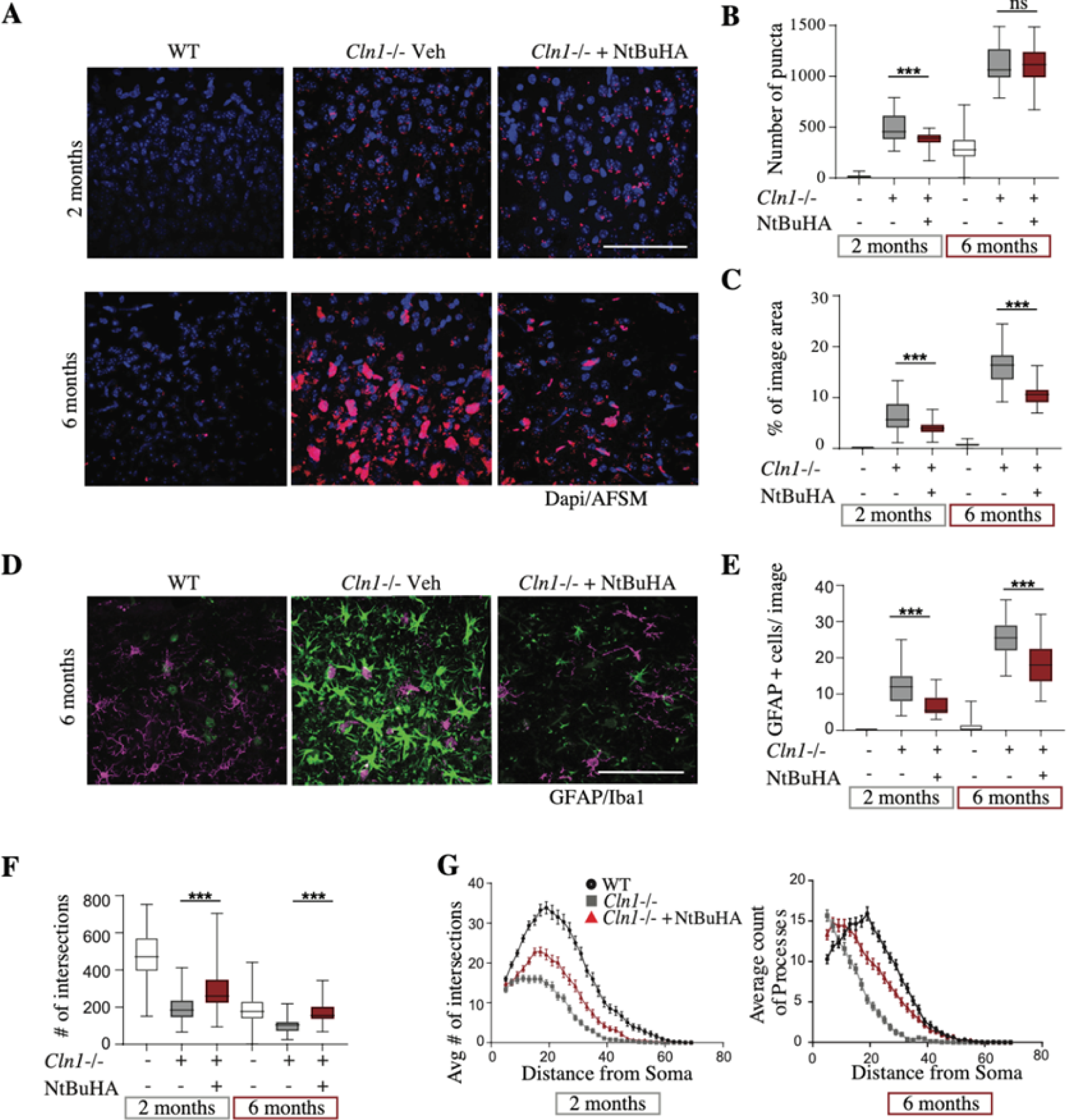
Reduction of AFSM Accumulation and Neuroinflammation in Cln1-/- Mice Following NtBuHA Treatment. **(A)** showcases representative micrographs depicting AFSM accumulation in the visual cortex of Wild Type (WT), untreated *Cln1*-/-, and *Cln1*-/- mice treated with 4mM NtBuHA, at 6 months of age (P180), Scale=100_μ_m. Quantification of **(B)** counts of AFSM puncta, 2 months: *Cln1*^−/−^ = 500.2 particles, n= 30; *Cln1*^−/−^ 1mM NtBuHA = 383.6 particles, n=31, P=0.0018, Dunnett’s multiple comparisons test. **(C)** the percentage of area covered by AFSM puncta in cortical sections, comparing WT, untreated *Cln1^−/−^*, and NtBuHA-treated *Cln1*^−/−^ mice at 2 and 6 months of age, 2 months: *Cln1*^−/−^ = 6.409, n= 30; *Cln1*^−/−^ 1mM NtBuHA = 3.933, n=31, P=0.0013; Dunnett’s multiple comparisons test; 6 months: *Cln1*^−/−^ = 15.97, n=39; *Cln1*^−/−^ 4mM NtBuHA = 10.77, n=43, P<0.0001, Tukey’s multiple comparisons test. **(D)** Representative images GFAP (Green) and Iba1(Magenta) immunostaining in the same 3 groups as (A) at 6 months of age. Scale=100_μ_m. **(E)** Quantification of GFAP cell count per image in WT, *Cln1*^−/−^, and NtBuHA-treated *Cln1*^−/−^ cortex at 2 and 6 months of age, 2 months: *Cln1*^−/−^ =12.19 cells, n=31; *Cln1*^−/−^ 1mM NtBuHA = 4.567 cells, n=31, P<0.0001, Tukey’s multiple comparisons test; 6 months: *Cln1*^-/*-*^ = 25.53 cells, n=34; Cln1^−/−^ 4mM NtBuHA = 18.06 cells, n=33, P<0.0001, Tukey’s multiple comparisons test. **(F)** Quantification of total microglial arborizations. 2 months: *Cln1*^−/−^ = 192.1 arborizations, n=54; *Cln1*^−/−^ 1mM NtBuHA = 292.4 arborizations, n=52, P<0.0001, Tukey’s multiple comparisons. 6 months: *Cln1*^−/−^ = 102.5 arborizations, n=60; *Cln1*^−/−^ 4mM NtBuHA = 167.8 arborizations, n=60, P<0.0001, Tukey’s multiple comparisons**. (G)** Quantification of the number of arbors as a function of the distance from the cell soma using Scholl analysis WT, *Cln1*^−/−^, and NtBuHA-treated *Cln1*^−/−^ cortex at 2 months (left panel) and 6 months (right panel). Data are represented as mean ± SEM. Data representation for B, C, E and F is in the format of box-and-whisker plots, indicating minimum to maximum values with a line representing the mean.

In CLN1, progressive neuroinflammation can be characterized by increased numbers of glial fibrillary acidic protein (GFAP)-expressing astrocytes [36–38]. NtBuHA treatment has been shown to reduce GFAP-reactive cells in *Cln1*^-/*-*^ mice at 6 months [26]. Building on these findings, we assessed the impact of chronic NtBuHA supplementation by quantifying GFAP-positive cells, and the area covered by GFAP immunolabeling in layer 2/3 of S1 and V1 in 2-and 6-month-old *Cln1*^-/*-*^ mice (**Fig. 2D-E, Supplemental Fig. 2**). A significant elevation in astrogliosis was noted between 2 and 6 months in *Cln1*^-/*-*^, aligning with disease progression (**Fig. 2E**). NtBuHA treatment significantly reduced astrogliosis both at 2 months and 6 months, with decreased GFAP-positive cell counts (**Fig. 2E**) and reduced area occupied by GFAP+ cells (**Supplementary Fig. 2C**).

An additional indicator of neuroinflammation in CLN1 that intensifies with disease progression is microglial activation, detectable by the adoption of a pro-inflammatory morphology [39–41]. Morphological analysis of ionized calcium-binding adaptor molecule 1 (Iba1)-positive microglia in S1 and V1 revealed that *Cln1*^-/*-*^ mice receiving NtBuHA exhibited an increased number of microglial process intersections and elongated average process length compared to untreated controls at both 2 and 6 months (**Fig. 2F-G**).

### 3.3 NtBuHA reduces synaptic calcium event frequency in Cln1^−/−^ neurons

Recent studies from our lab demonstrated that *Cln1*^-/*-*^ neurons exhibit aberrant postsynaptic calcium dynamics that underpin neuronal hyperexcitability *in vitro* and correspond to synchronous cortical neuron firing *in vivo* [29]. Therefore, we conducted a live cell imaging assay [28,29] in WT, *Cln1*^-/*-*^, and NtBuHA-treated *Cln1*^-/*-*^ primary cortical neurons to analyze the properties and dynamics of postsynaptic calcium influx at individual synapses (**Fig. 3A, 3B**). We first measured the amplitude of spontaneous calcium transients at dendritic boutons. Consistent with a previous report [28], spontaneous calcium transients were larger in *Cln1*^-/*-*^ neurons than in WT cells, whereas NtBuHA-treated neurons demonstrated a surprising increase in calcium transient amplitude compared to untreated *Cln1*^-/*-*^ neurons (**Supplemental Fig. 3A**). To test whether the amplitude of the most prominent synaptic calcium events in *Cln1*^-/*-*^ neurons is affected by NtBuHA treatment, we considered the 200 highest amplitude calcium events from each group (**Fig. 3C; Supplemental Fig. 3B**). Using this metric, we again found that the largest spontaneous calcium transients were of greater amplitude in *Cln1*^-/*-*^ neurons than in WT cells, while there was no difference between untreated and NtBuHA-treated *Cln1*^-/*-*^ cells (**Fig. 3C Supplemental Fig. 3B**). Given this finding, we next considered the rate of spontaneous calcium events across each group. Here, we found that NtBuHA treatment had a marked effect on calcium event frequency. Specifically, while *Cln1*^-/*-*^ neurons demonstrated a greater instantaneous event frequency compared to WT cells, exaggerated event frequency was normalized following NtBuHA treatment (**Fig. 3D, 3E)**.

**Figure 3:**
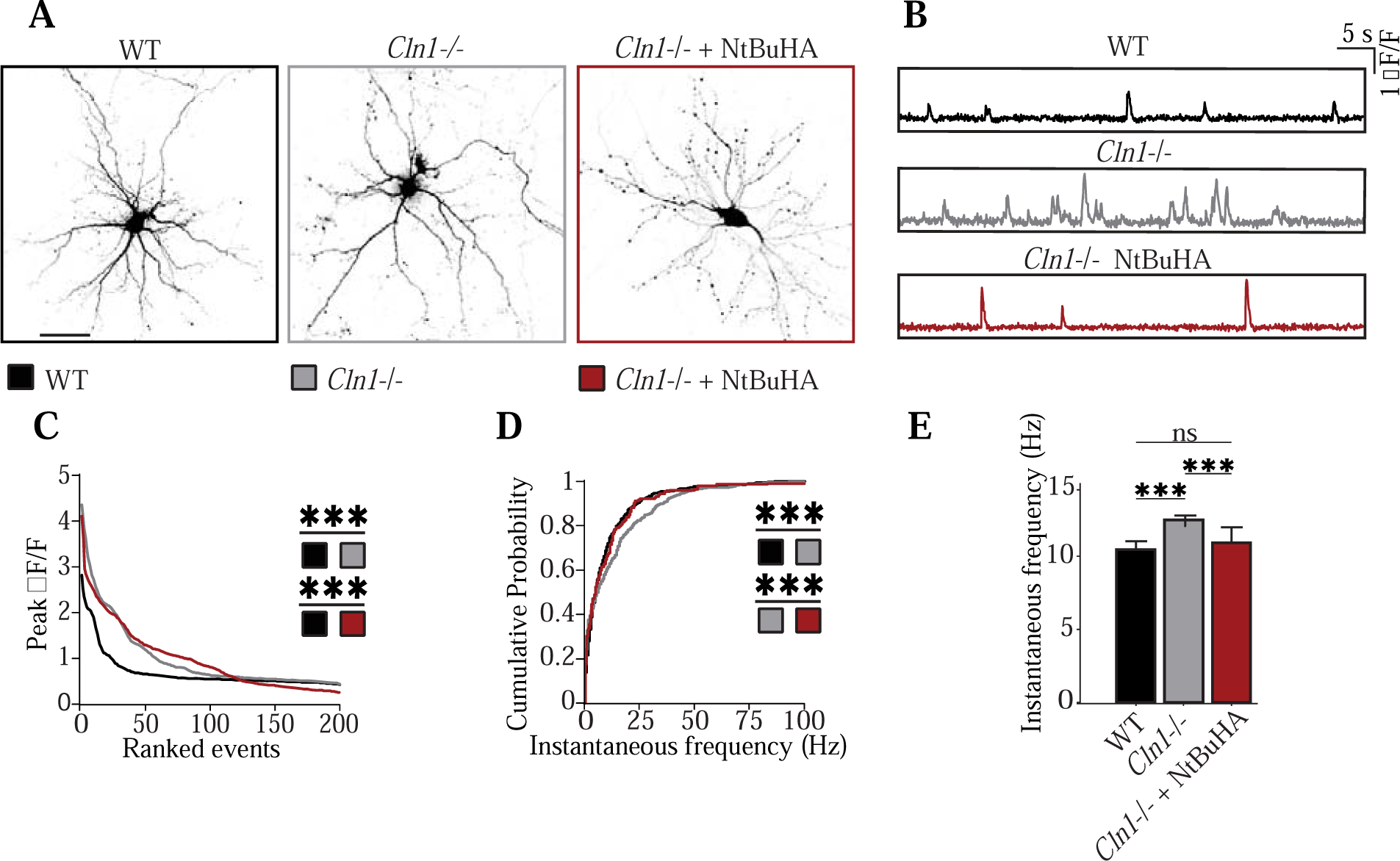
Normalization of Synaptic Calcium Event Frequency via NtBuHA. (**A**) Example images of WT, *Cln1*^−/−^, and NtBuHA-treated *Cln1*^−/−^ neurons. Scale bar = 50 um. (**B**) Example spontaneous calcium transients extracted from WT, *Cln1*^−/−^, and NtBuHA-treated *Cln1*^−/−^ neurons. (**C**) The amplitudes of the 200 largest events from each group are sorted in order. Mean amplitude values are as follows *Cln1^−/−^* = 0.92 ± 0.05, dF/F; WT = 0.60 ± 0.03 dF/F, n = 200; P < 0.001, rank sum test; WT = 0.60 ± 0.03 dF/F, n=200; *Cln1^−/−^ + NtBuHA =* 0.93 ± 0.05, dF/F, n = 200; P <0.001 rank sum test. (**D-E**) Cumulative distributions of the instantaneous frequency of all detected calcium events from each group, *Cln1*^−/−^ = 12.52± 0.65 Hz, n = 596; WT = 10.47 ± 0.52 Hz, n = 740; P < 0.001 rank sum test; *Cln1* ^−/−^ + NtBuHA = 10.95 ± 1.04 Hz, n = 191; P < 0.001, comparison to mutant untreated cells; rank sum test.

### 3.4 Impact of NtBuHA on Seizure Dynamics and Motor Function in Cln1-/- Mice

Disease-modifying treatments for CLN1 Disease have historically generated mixed results on improving seizures in CLN1 mouse models [37,42]. We conducted EEG monitoring of epileptiform activity (represented by high amplitude spike trains) in WT and *Cln1*^-/*-*^ animals treated long-term (from 1-to 7-months of age) with either NtBuHA (4mM) or vehicle. For a subset of EEG recordings, time-stamped video recordings confirmed dynamic spike detection thresholding and were consistent with characteristic behavioral phenotypes of convulsant seizures recently reported in *Cln1*^−/−^ mice (**Supplemental Video 1-2**) [42]. Crucially, treatment with NtBuHA diminished the number of spike trains (**Fig. 4A; Supplemental Fig. 4**), as well as the total and average spike train duration in *Cln1*^−/−^ mice (**Fig.4B, 4C**). Intriguingly, we found that NtBuHA-treated *Cln1^−/−^* mice had increased spike frequency, albeit this effect did not reach statistical significance (**Figure 3D**). This seemingly paradoxical trend is consistent with human clinical studies showing diverse effects of anti-seizure medications on epileptic event propagation [11,12,43]. Our findings provide the first evidence for an effective treatment strategy that reduces seizures in *Cln1^−/−^*mice.

**Figure 4.**
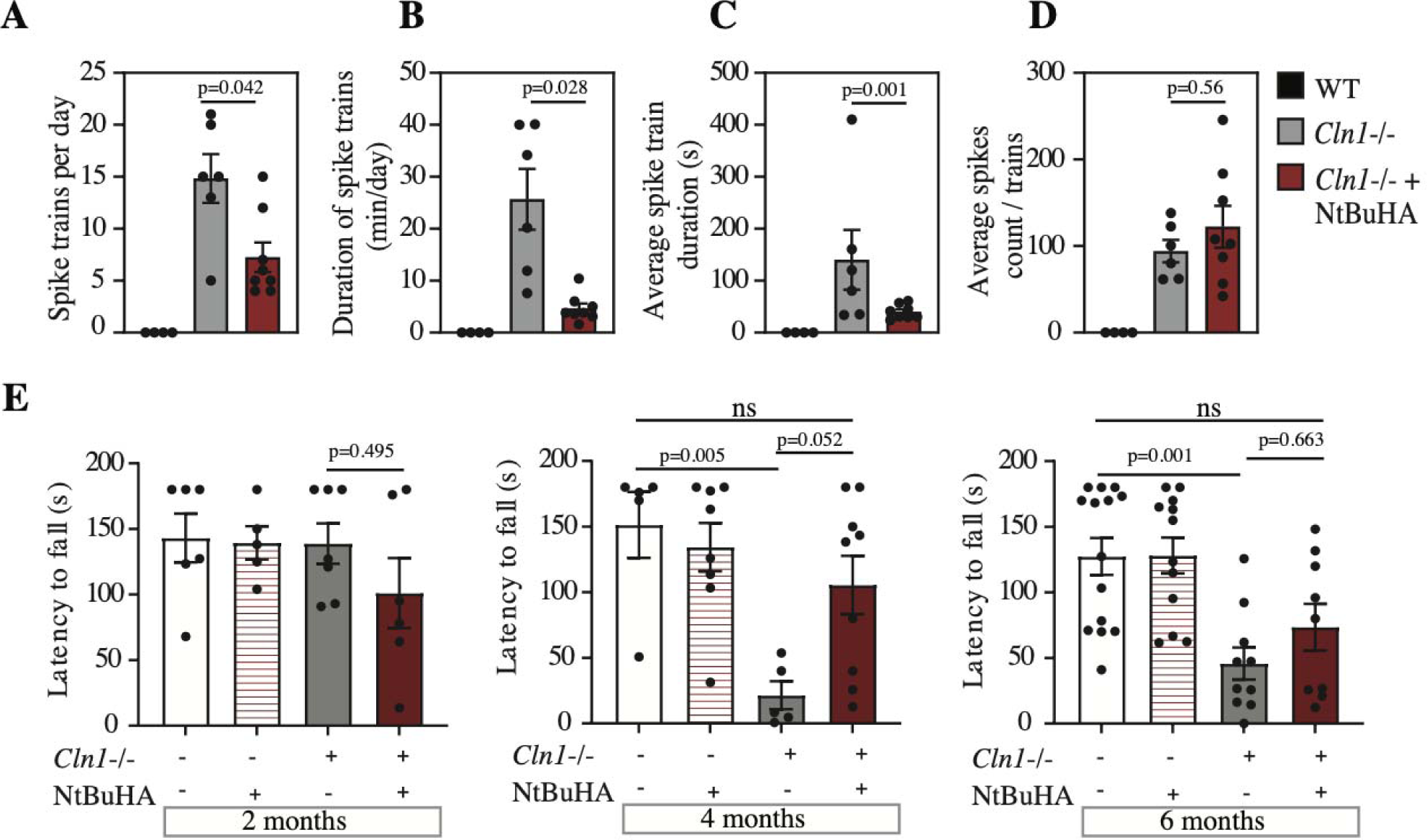
Efficacy of NtBuHA in Modulating Seizure Activity and Motor Coordination in Cln1-/- Mice. Quantification of **(A)** total number of spike trains, *Cln1*^−/−^ = 12.7 spikes, n= 6; *Cln1*^−/−^ 4mM NtBuHA = 6.07 spikes, n= 8, P=0.0072, Tukey’s multiple comparisons test **(B)** cumulative duration of spike trains, *Cln1*^−/−^ =23.95 minutes, n= 6; *Cln1*^−/−^ 4mM NtBuHA = 3.739 minutes, n=8, P=0.0037, Tukey’s multiple comparisons test **(C)** average duration of spike trains, *Cln1*^−/−^ = 140.2 minutes, n= 6; *Cln1*^−/−^ 4mM NtBuHA = 40.06, n= 8, P=0.001, Tukey’s multiple comparisons test and **(D)** average spikes per train, compared across Wild Type (WT), *Cln1*^−/−^, and NtBuHA-treated (4mM) mice, using a spike detection threshold set at 5x baseline (Mann-Whitney test, p-values indicated), *Cln1*^−/−^ = 94.15 spikes, n=6; *Cln1*^−/−^ 4mM NtBuHA = 122.3 spikes, n=8, P=0.5634, Tukey’s multiple comparisons test. **(E)** Rotarod performance tests at 2, 4, and 6 months of age for WT, NtBuHA-treated WT, *Cln1*^−/−^, and NtBuHA-treated *Cln1*^−/−^ mice (n=5-9 mice/group) reveal motor coordination, analyzed using ANOVA followed by Tukey’s test (p-values shown), **2 Months;** WT= 143.1, seconds, n= 6; *Cln1*^−/−^ = 138.9, seconds, n=7, P=0.9985, WT= 143.1, seconds, n= 6; *Cln1*^−/−^ 1mM NtBuHA = 101.1, seconds, n= 6, P=0.4368, *Cln1*^−/−^ = 138.9, seconds, n=7; *Cln1*^−/−^ 1mM NtBuHA = 101.1, seconds, n= 6, P=0.4954, Tukey’s multiple comparisons test, **4 Months;** WT= 151.3, seconds, n= 5; *Cln1*^−/−^ = 21.46, seconds, n= 7, P=0.0055, WT= 151.3, seconds, n= 5; *Cln1*^−/−^ 4mM NtBuHA = 105.5, seconds, n=9, P=0.4572, Tukey’s multiple comparisons test, **6 Months;** WT=127.3, seconds, n=15; *Cln1*^−/−^ = 45.71, seconds, n= 9, P=0.0012, WT=127.3, seconds, n=15; *Cln1*^−/−^ 4mM NtBuHA = 73.51, n=9, P=0.06, Tukey’s multiple comparisons test.

Finally, we assessed motor function using a fixed speed rotarod assay. Rotarod testing at 2 months of age showed no difference in the average latency to fall between WT, *Cln1*^-/*-*^, and NtBuHA-treated *Cln1*^-/*-*^ mice (**Fig.4E, *left*)**. At 4 and 6 months of age, untreated *Cln1*^-/*-*^ animals demonstrated a significantly shorter latency to fall compared to WT counterparts. In contrast, *Cln1*^-/*-*^ mice treated with NtBuHA (P30-P180) had no difference in latency to fall compared to WT (**Fig.4E, *middle and right*)**. Indeed, the behavioral performance of NtBuHA-treated *Cln1*^-/*-*^ animals at 4 months approached, but did not quite reach, a statistical improvement compared to untreated mice (**Fig.4E, *middle*)**.

## 4. Discussion

This study investigated the therapeutic effects of orally administered PPT1-mimetic small molecule, NtBuHA (CIRC827), in *Cln1^−/−^*mice. The results demonstrate that NtBuHA treatment decreased histological features of neuroinflammation, reduced AFSM accumulation, improved motor function, corrected synaptic calcium dynamics, and decreased the number and duration of seizures. These findings suggest a mechanism for how NtBuHA could be reducing seizure severity and indicate NtBuHA supplementation improves multiple layers of pathology in *Cln1^−/−^* mice.

### 4.1 Potential mechanisms for reduction of seizure with NtBuHA treatment

The global neuroinflammatory phenotype observed in *Cln1^−/−^*mice was robustly suppressed by NtBuHA treatment. While only 30 days of treatment (P30-P60) was sufficient to reduce gliosis, chronic treatment from one to six months blunted late-stage neuroinflammation as well. These data corroborate previous findings using NtBuHA, which demonstrate similar improvements to markers of neuroinflammation in four-and six-month *Cln1*^−/−^ mice [44]. Here, we provide additional evidence that microglial activation is also reduced by PPT1-mimetic treatment at six months [26]. While the presence of gliosis serves as a histopathological correlate of disease progression, it has also been causally implicated in several CLN1 disease processes, including the generation of seizures [45–47]. Further, microglial activation is associated with seizure onset in a CLN1 mouse model [40]. Therefore, one potential interpretation is that NtBuHA treatment suppressed seizures via a reduction in neuroinflammation [40,42,44,48].

However, countervailing evidence for this interpretation comes from experiments employing chronic cannabidiol treatment in *Cln1^−/−^*mice [42], which suggest that neuroinflammatory severity and the emergence of seizures may be decoupled. That is, whereas cannabidiol supplementation successfully reduced microglial neuroinflammation, no parallel improvement in epileptic activity was observed [42]. Thus, while we suspect that diminishing neuroinflammation had a positive effect on seizure activity in NtBuHA-treated mice, the anti-seizure effects of the drug may also be driven by other mechanisms. Indeed, our findings allude to a second potential mechanism by which NtBuHA treatment reduced seizures in *Cln1^−/−^*mice.

A growing body of evidence demonstrates that disrupted calcium dynamics in *Cln1^−/−^* neurons correlates with circuit-wide alterations to cortical neuron firing patterns [28,29]. Here, we demonstrate that chronic supplementation of NtBuHA in the medium of *Cln1*^−/−^ primary cortical neurons corrected the increased frequency of postsynaptic calcium transients compared to untreated *Cln1*^−/−^ neurons. Supporting this finding, recent work [48] shows that NtBuHA treatment impacts AMPA receptor transmission in hippocampal slices, although the effect was a reduction in the amplitude, rather than the frequency, of spontaneous postsynaptic events [48]. This discrepancy is reconciled by methodological differences. Namely, the study by Xia *et al.*, used an acute (30 minute) NtBuHA incubation and made physiological recordings of spontaneous postsynaptic events, which are AMPA receptor mediated, whereas we used a chronic treatment paradigm and imaged postsynaptic calcium transients, which involve both calcium permeable AMPA receptors and NMDA receptors [28,29]. These are crucial differences, considering Xia and colleagues noted a reduction in AMPA, but not NMDA receptor palmitoylation following acute NtBuHA treatment [48], while our previous data show that chronic palmitoylation inhibitor treatment can impact NMDA receptor palmitoylation [28]. Furthermore, we studied NtBuHA effects in *Cln1*^−/−^ neurons, which demonstrate dysregulation of both glutamate receptor subtypes [28,29]. Taken together, it appears that NtBuHA treatment duration selectively modulates glutamate receptor subtypes and their trafficking. Specifically, whereas acute treatment influences the short-term trafficking of AMPA receptors, chronic treatment likely affects both AMPA and NMDA receptor dynamics.

Nevertheless, a holistic view of the evidence indicates that NtBuHA modulates synaptic activity and calcium entry through glutamate receptors. As abnormal neuronal calcium dynamics are implicated in epileptogenesis [49–51], this finding highlights the intriguing possibility that NtBuHA supplementation in *Cln1*^−/−^ mice suppressed seizures through correcting synaptic calcium dynamics [51–53]. However, more experiments are required to determine whether the NtBuHA-mediated correction of synaptic transient frequency *in vitro* can be extrapolated to intact animals and whether such an effect is responsible for the alleviation of seizures observed in treated *Cln1*^−/−^ mice. Nevertheless, the reduction of seizure event frequency and duration in *Cln1^−/−^*mice is clinically significant, motivating further understanding of the likely broad mechanism of action of NtBuHA.

### 4.2 Current therapeutic landscape for CLN1 and the translational potential of NtBuHA

The current clinical standard for CLN1 treatment is palliative care, as there are no approved treatment options that target the underlying cause of the disease. The long-term goal is therefore to approve NtBuHA for use in CLN1 and provide clinicians the option of incorporating an orally administered, disease-treating small molecule into a patient’s treatment plan.

Beyond the advances of NtBuHA as a small molecule-based treatment option for CLN1, significant advances have also been made in the development of both gene therapies and enzyme replacement therapies (ERTs). Viral-mediated gene therapies have shown promise for the treatment of CLN1 across several studies and treatment paradigms [20–22,52,53], and several important recent advancements in viral vector technology are working toward realizing the safe systemic delivery of a blood-brain barrier (BBB)-penetrant virus to achieve robust, widespread transduction of CNS cell-types [55,56]. Further, ERT has proven effective in *Cln1^−/−^*mice [37,57,58], with the most promising results coming from studies utilizing intracerebroventricular or intrathecal administration that allows intact enzyme to bypass the BBB [37,58]. However, the clinical application of both gene therapy and ERT approaches remains limited by a need to optimize the delivery (i.e., achieve widespread expression of the functional enzyme) and suppress potential adverse events [54].

Therefore, small molecule-based therapeutic strategies, either in isolation or in combination with gene therapy or ERT approaches, may offer a strategic benefit to treating CLN1. Many small molecules such as NtBuHA have good oral bioavailability and can cross the BBB, and thus may be orally administered. In addition to the practical benefits of oral administration, systemic delivery via this route allows for widespread distribution of drug to impact both the CNS and periphery. This feature may become increasingly important considering the growing evidence that CLN1 Batten Disease affects the spinal cord and enteric nervous system [55,56]. Further, with brain penetrant small molecules, distribution of the small molecule through the brain may supplement regions that may be more difficult to reach by gene therapies or ERTs through intrathecal or intraventricular injections [26,57].

Certainly, there remains an unmet need for a safe and effective curative therapy for CLN1 disease. The recent impactful study [37] demonstrating the efficacy of ERT in multiple animal models is promising. Here, we provide evidence for an additional near-term candidate for treating CLN1. Ongoing work includes further characterization of the dose-dependent pharmacodynamic and pharmacokinetic behavior of NtBuHA, as well as a Good Laboratory Practices (GLP)-grade toxicology and safety pharmacology program in preparation for a First-in-Human trial. Clear next steps are to evaluate this formulation of NtBuHA (CIRC827) in a larger preclinical context, including larger animal models [24,37], in addition to testing its efficacy in combination with gene therapy and/or ERT approaches.

### 4.3 Therapeutic potential for NtBuHA beyond CLN1

We demonstrate herein that NtBuHA has manifold effects on neuronal and glial function in vivo. While an immediate goal is to optimize and employ NtBuHA to alleviate the suffering of CLN1 patients, our findings also point to a broader therapeutic potential for this small molecule. Firstly, lysosomal disruption is a characteristic of most common neurodegenerative diseases [57,58] including Alzheimer’s disease [59], and NtBuHA reproducibly decreases lysosomal waste accumulation and improves lysosomal function [26,60]. Similarly, neuroinflammation is a common feature of many neurological conditions that is alleviated by NtBuHA treatment [61]. Finally, while protein palmitoylation and depalmitoylation must be balanced for proper cellular function, this is particularly true for neurons. Indeed, synaptic proteins are significantly enriched for palmitoylation sites compared to the broader proteome [62], suggesting an outsized requirement for regulation of protein palmitoylation in the brain. It follows that dysregulated protein palmitoylation is associated with several neurological conditions, like Alzheimer’s disease, where palmitoylation of the enzymes that process amyloid precursor protein affect disease progression [63]. This presents the intriguing possibility that therapeutics targeting protein palmitoylation will have particularly strong effects in the CNS, where treatments have a very high failure rate. Taken together, NtBuHA targets several shared features of common neurological and neurodegenerative diseases, suggesting a broad therapeutic potential for NtBuHA beyond CLN1.

## Author contributions

Zach Fyke: investigation, data curation, writing-original draft and editing, visualization. Rachel Johansson: writing-original draft and editing. Joseph D. Zak: formal analysis, writing-original draft and editing. Anna I. Scott: writing-original draft and editing, funding acquisition. Devin Wiley: funding acquisition, conceptualization, writing-original draft and editing. Daniel Chelsky: funding acquisition, project administration, conceptualization, writing-original draft and editing. Kevin P. Koster: supervision, methodology, writing-original draft and editing. Nader Al-Nakouzi: project administration, conceptualization, visualization, writing-original draft and editing. Akira Yoshii: supervision, resources, methodology, conceptualization.

## Acknowledgments

The authors would like to thank Kamal Sharma and Lisa Hoffman for providing EEG recording hardware and training. We would also like to thank various past Yoshii lab members for helpful conversations and support. Additionally, this project is dedicated to Akira Yoshii, who sadly lost his battle with cancer during the course of this project. As a true physician-scientist, Akira’s goal was to bridge the gap between bench & bedside. He hoped to accomplish this feat by conducting translational research in his lab, then administering effective treatments to children suffering from pediatric neurological diseases. We hope this study, along with others published posthumously, will in part fulfil his life’s purpose.

---

**Supplemental Figure 1:**
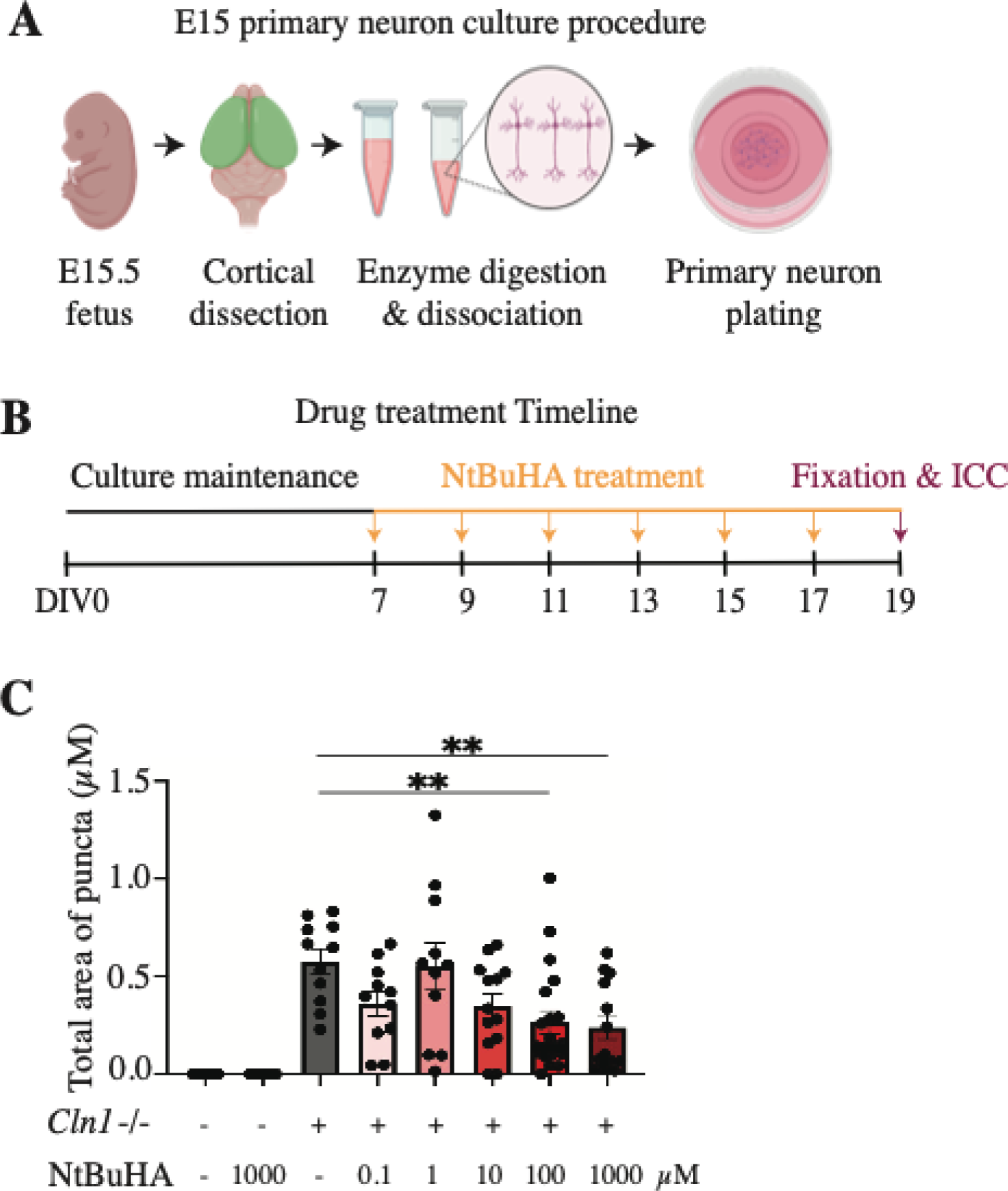
(**A**) Schematic of primary cortical neuron culture. (**B**) Timeline of primary cortical neuron treatment with NtBuHA before ICC. (**C**) Graph representing the total area of AFSM puncta, *Cln1^−/−^* vehicle-treated = 0.5787 total area of puncta, n=29; comparison to *Cln1^−/−^* 100mM NtBuHA = 0.266 total area of puncta, n=35, P=0.0089; comparison to *Cln1^−/−^*1mM NtBuHA = 0.239 total area of puncta, P=0.0095, Tukey’s multiple comparisons test.

**Supplemental Figure 2:**
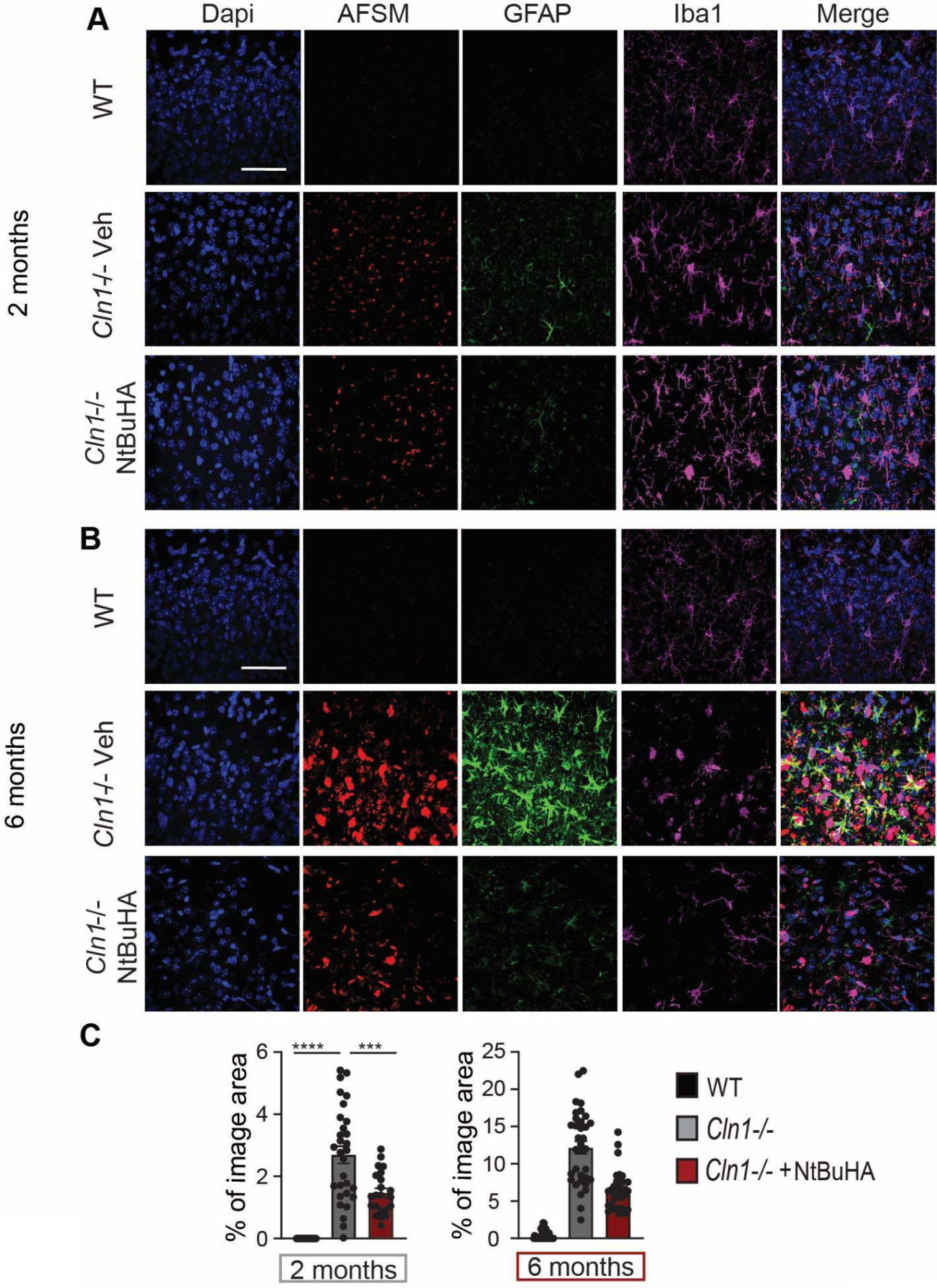
(**A**) Example images of V1 coronal sections from WT, Cln1^−/−^, and NtBuHA-treated Cln1^−/−^ mice at 2 months of age, and (**B**) at 6 months of age. Scale bar = 50 um. (**C**) Percent of image area for GFAP stained brain sections at 2 and 6 months respectively, 2 months; *Cln1^−/−^*=2.835 percent area, n=31; *Cln1^−/−^* 1mM CIRC827=0.9908 percent area, n=31, P<0.0001, Tukey’s multiple comparisons test; 6 months; *Cln1^−/−^* =12.18 percent area, n=34; *Cln1^−/−^* 4mM CIRC827=6.418 percent area, n=33, P<0.0001, Tukey’s multiple comparisons test.

**Supplemental Figure 3:**
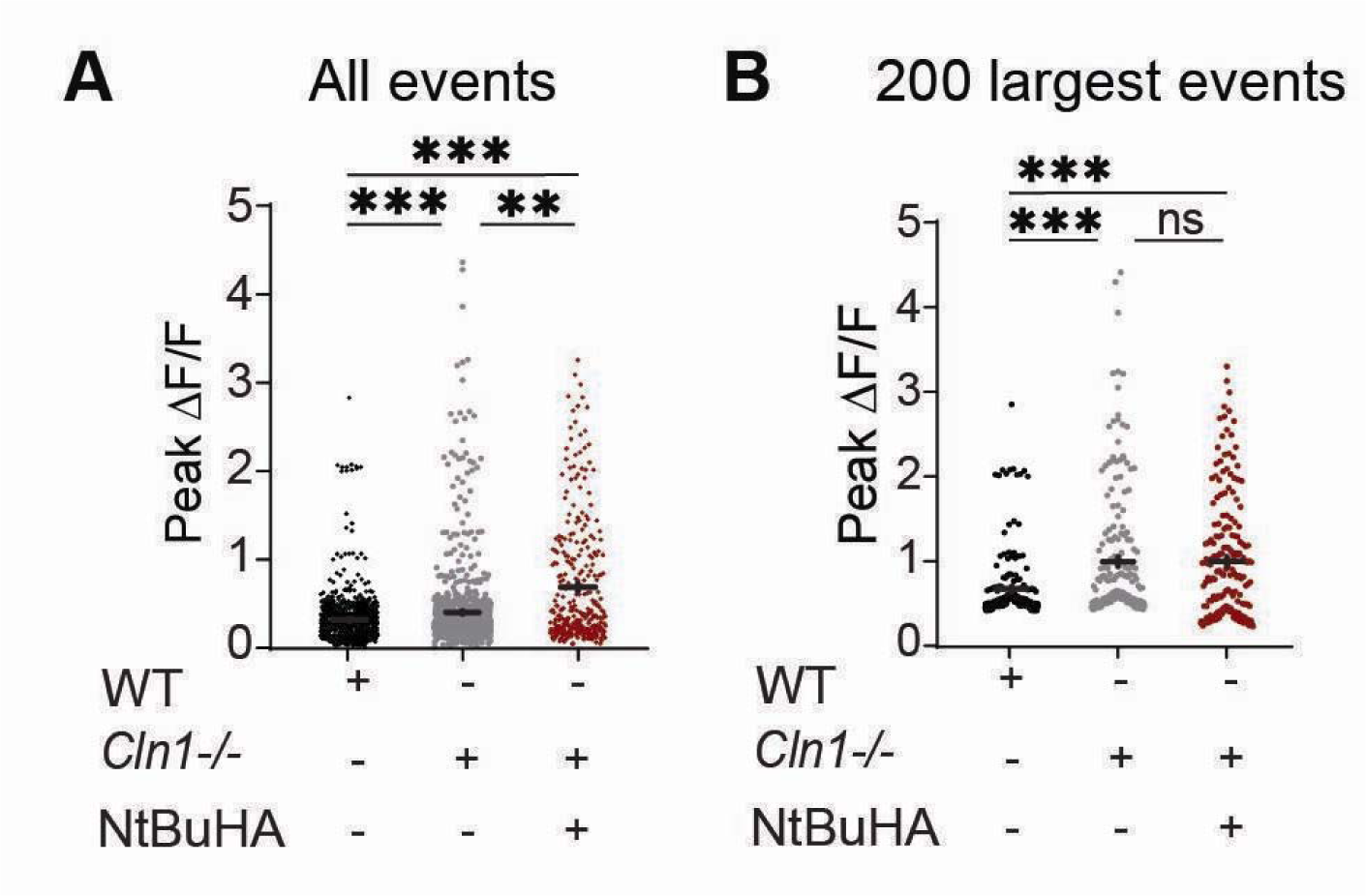
(**A**) Quantification of the calcium transient amplitude for all synaptic sites (represented by individual points on graph) from all neurons: *Cln1*^−/−^ = 0.36 ± 0.02, dF/F, n = 899; *Cln1*^−/−^ + NtBuHA = 0.64 ± 0.04, dF/F, n = 324; P < 0.001; WT = 0.28 ± 0.01 dF/F, n = 824; P < 0.001, Kolmogorov-Smirnov test. (**B**) Amplitudes of the 200 largest events: *Cln1^−/−^* = 0.92 ± 0.05, dF/F; WT = 0.60 ± 0.03 dF/F, n = 200; P < 0.001, rank sum test; WT = 0.60 ± 0.03 dF/F, n = 200; *Cln1^−/−^ + NtBuHA =* 0.93 ± 0.05, dF/F, n = 200; P <0.001 rank sum test; *Cln1^−/−^ + NtBuHA =* 0.93 ± 0.05, dF/F, n = 200 P = 0.25, comparison to *Cln1^−/−^* untreated cells; rank sum test.

**Supplemental Figure 4.**
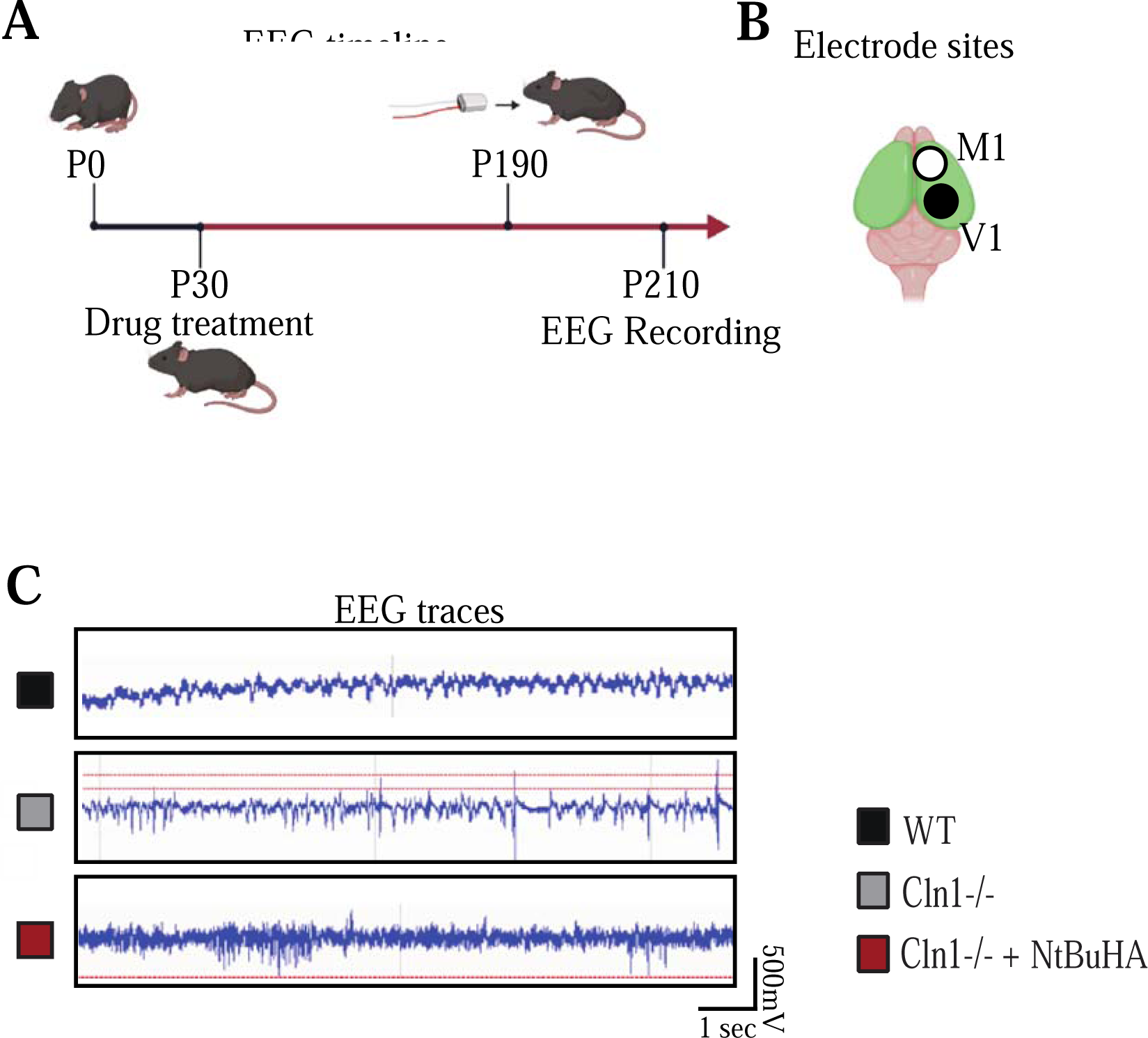
(**A**) Timeline of NtBuHA treatment and ETA-F10 telemetry implantation. (**B**) Graphical representation of positive (White) and negative (Black) subdural electrode implantation sites. (**C**) Representative traces of EEG signals from 7-month-old WT, *Cln1^−/−^,* and *Cln1^−/−^* NtBuHA-treated mice.

**Supplemental Video 1. Simultaneous video tracking and EEG recording of non-convulsant seizures.** (**A**) Video (left) and time-locked EEG recording (right) from a Vehicle-treated Cln1^−/−^ mouse at seven months demonstrating severe hypertonia of the tail, alongside behavior associated with high-frequency, high-amplitude spikes in the EEG trace.

**Supplemental Video 2. Simultaneous video tracking and EEG recording of convulsant seizures.** (**B**) Video (left) and time-locked EEG recording (right) from a NtBuHA-treated *Cln1*^−/−^ mouse at seven months demonstrating the correlation between a convulsive seizure and high-frequency, high-amplitude spikes in the EEG trace. Note the shortened seizure duration in the NtBuHA-treated animal.

## Resources Table

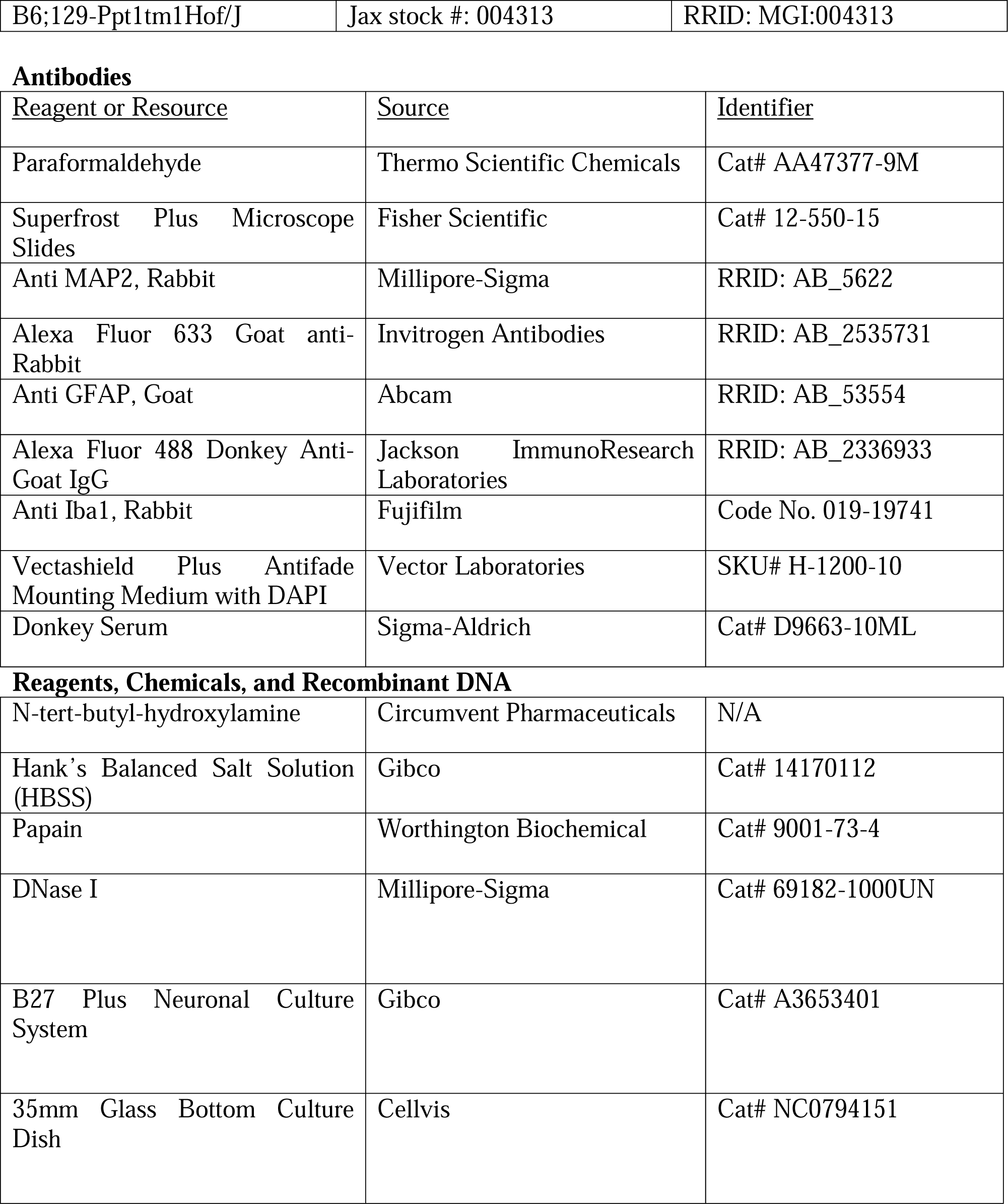
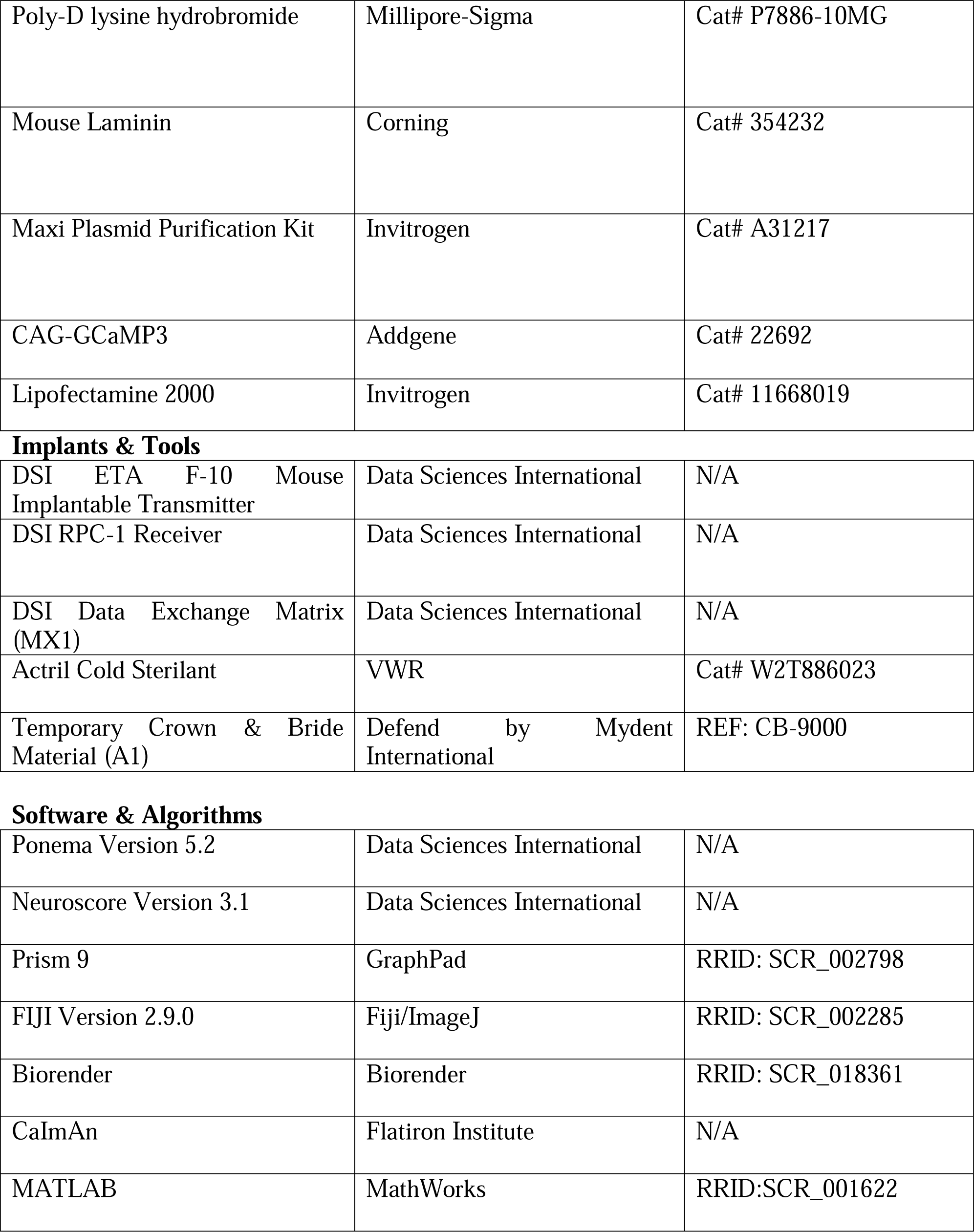

